# Subcortical coding of predictable and unsupervised sound-context associations

**DOI:** 10.1101/2022.10.06.511202

**Authors:** Chi Chen, Hugo Cruces-Solís, Alexandra Ertman, Livia de Hoz

## Abstract

Our environment is made of a myriad of stimuli present in combinations often patterned in predictable ways. For example, there is a strong association between where we are and the sounds we hear. Like so many environmental patterns, sound-context associations are learned implicitly, in an unsupervised manner, and are highly informative and predictive of normality. Yet, we know little about where and how unsupervised sound-context associations are coded in the brain. Here we measured plasticity in the auditory midbrain of mice living over days in a naturalistic environment designed to present sound-context associations with different degrees of predictability. Plasticity in the auditory midbrain, a hub of auditory input and multimodal feedback, developed over days and reflected learning of contextual information in a manner that depended on the predictability of the sound-context association and not on reinforcement. Plasticity took the form of broad frequency shifts in tuning in auditory midbrain neurons. These shifts were paralleled by an increase in response gain and correlated with an increase in neuronal frequency discrimination. Thus, the auditory midbrain codes for unsupervised predictable sound-context associations, revealing a subcortical engagement in the detection of contextual sounds. This detection might facilitate the processing of behaviorally relevant foreground information described to occur in cortical auditory structures.

## Introduction

Our soundscape is laden with patterned information, much of which can be learned and then predicted. This statistical learning (Reber, 1967) is evident in the strong association that our brain makes between places and sounds. Indeed, as animals move through their environment, they implicitly, without reinforcement, learn and can often predict sound-context pairings. As we enter an environment, being able to quickly assess whether these sound-context pairings match our prediction can be essential for survival, yet we know little about the circuits and mechanisms underlying their representation. In this study, we investigated sound coding in the subcortical auditory midbrain of mice living in an enriched environment with different degrees of predictability in the sound-context pairings. The auditory midbrain (inferior colliculus) was chosen because it is a hub of sensory input and multimodal feedback (Nakamoto et al., 2013; Yudintsev et al., 2021), known to be involved in the processing of soundscapes (Cruces-Solís et al., 2018).

Reinforced learning of sound-reinforcement pairings leads to plasticity in auditory structures. It takes the form of frequency-specific tuning shifts towards the reinforced tone in the auditory cortex (Bieszczad and Weinberger, 2010; Blake et al., 2002; Letzkus et al., 2011) and auditory midbrain (Gao and Suga, 2000; Yan et al., 2005; Zhang and Suga, 2005). Evidence for the detection and coding of unsupervised (non-reinforced) contextual sounds also exists. For example, prior exposure to the sounds in a room can improve speech understanding in listeners (Brandewie and Zahorik, 2018) and influence our perception of the world in a pre-attentive manner (McPherson and McDermott, 2020). Similarly, cortical auditory structures show noise-invariant responses, which implies knowledge of the structure of the background noise (Khalighinejad et al., 2019; Mesgarani et al., 2014; Rabinowitz et al., 2012). Our own work has determined that exposure over days to a sound that was predictably played whenever the animal entered and stayed in a context (predictable sound-context association) leads to broad shifts in tuning in the inferior colliculus (IC) (Cruces-Solís et al., 2018). Here we aimed to further study the role of different contextual factors, specifically the presence of a reinforcer, the predictability of the sound onset, and the sound exposure time, in the development of associative plasticity in the auditory midbrain.

## Results

### 1. Contextual learning elicits IC plasticity independently of reinforcer

We first examined the role of reinforcement on the plasticity elicited by predictable sound-context associations in the auditory midbrain. We separately exposed three groups of mice in the Audiobox (Fig 1A), an enriched and automatic apparatus made of 2 main areas connected by corridors. Here, mice live in groups for days with little interaction with the experimenter. Each group had a different sound-context pairing experience in the sound corner (see Methods, Fig 1A). Importantly, none of the animals were conditioned and food and water were always *ad libitum*. Exposed mice, in the “predictable-W” (W: water) and “predictable-noW” groups (noW: no water; Fig 1A middle and right), heard 16 kHz tone pips for the duration of each visit to the sound corner. Thus, the act of entering the corner was predictive of sound onset. The control group (Fig 1A left) on the other hand, lived in the Audiobox for the same length of time but was not exposed to the 16kHz sound in the sound corner. Typically, when the water is in the corner, mice drink for about 60% of the visit time and in about 70% of the visits (Cruces-Solís et al., 2018; de Hoz and Nelken, 2014). Unlike the control and predictable-W groups, the “predictable-noW” group had no access to water in the corner and, instead, water bottles were in the food area, where no sound was ever played. This ensured that no association was made between the sound and the water. Then, to ascertain auditory midbrain plasticity, we acutely recorded from the left inferior colliculus (IC) of anesthetized animals that had been sound-exposed in the Audiobox for 6 to 12 days. For this experiment, we recorded multiunit activity using NeuroNexus multi-electrode arrays (NeuroNexus, MI, USA) with 16 sites linearly arranged and separated by 50 μm (Fig 1B). The electrode was dorsoventrally inserted in IC along the tonotopic axis and was placed in two consecutive steps to cover a range of depths between 0 and 1250 μm (see Methods; Fig 1B). To characterize frequency tuning, we presented sweeps of 30 ms long pure tones of different frequency-intensity combinations (see Methods) and built tuning curves for each recording site.

**Fig. 1:**
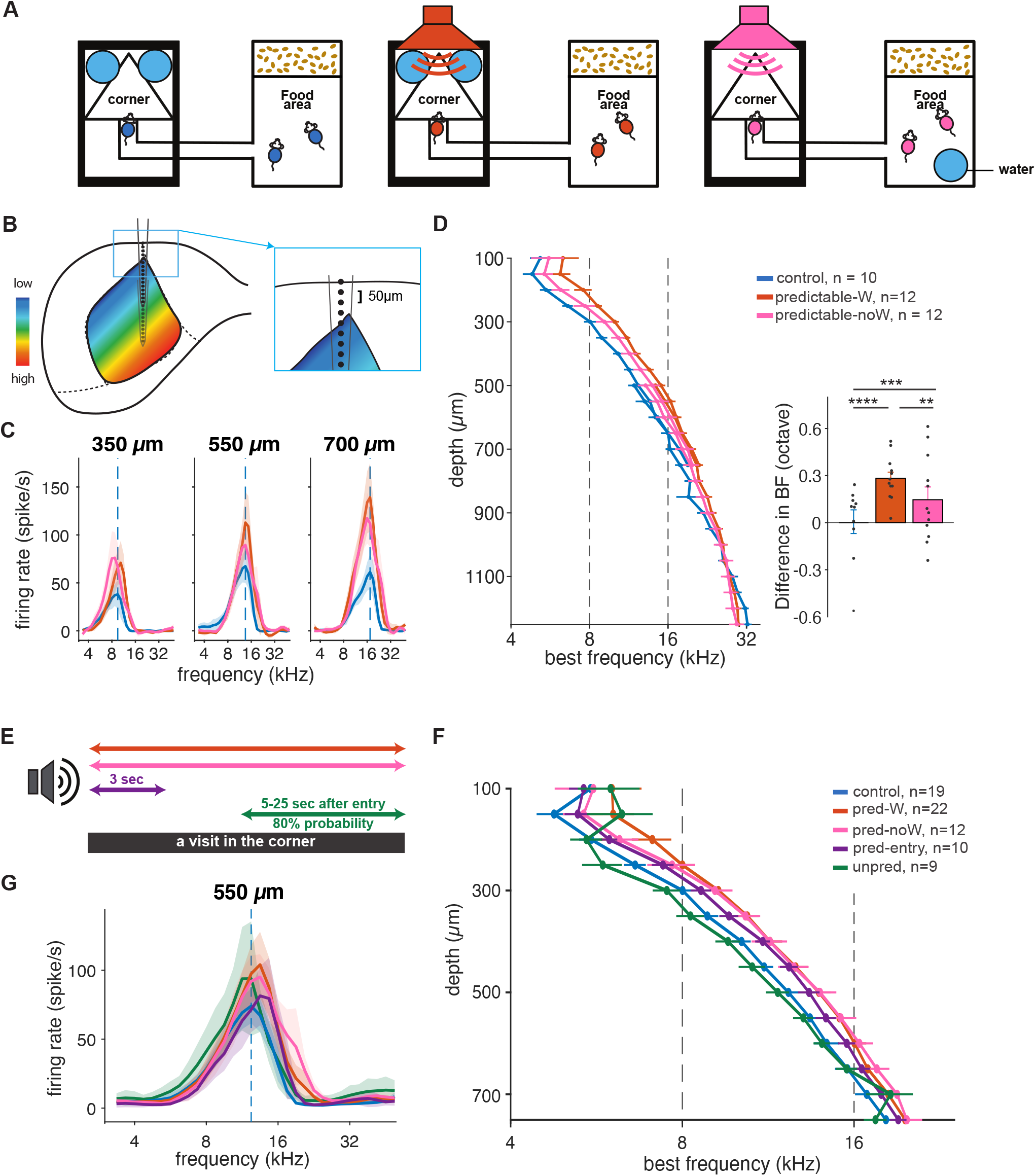
Exposure to predictable sound, independently of reinforcer, elicits IC plasticity. **A,** Schematic representation of the Audiobox design for the control (left), predictable (middle), and no-W (no-water) predictable exposure group (right), respectively. Food was *ad libitum* in the food area. Water was available in the corner (control and predictable group) or in the food area (no-W group). **B,** Schematic representation of recording using NeuroNexus 16-electrode arrays from two positions. Insert: Schematic representation of the positioning of the most superficial recording site, aligned to the dura. **C,** Average tuning curves of simultaneously recorded sound-evoked activity at 70 dB for the depth of 350, 550, and 700 μm in the IC of the control (blue), predictable (red), and no-W predictable (pink) group. **D**, Left: Average best frequency (BF) as a function of recording depths (two-way ANOVA, group *F*(2, 761) = 6.88, *p* = 0.001, depth *F*(25, 761) = 211.4, *p* = 0, interaction *F*(59, 761) = 0.85, *p* = 0.77). Right: Difference in BF relative to the average BF of the control group across the depths between 100 and 900 μm (two-way ANOVA, group *F*(2, 525) = 35.64, *p* = 0, depth *F*(16, 525) = 0.55, *p* = 0.92, interaction *F*(32, 525) = 0.52, *p* = 0.99. Corrected pair comparisons: \**p* < 0.05, \*\**p* < 0.01, \*\*\**p* < 0.001, \*\*\*\**p* < 0.0001). **E,** Schematic representation of the sound exposure protocol. **F,** Average BF for selected depths in the IC. **G,** Average tuning curves for the depth of 550 μm in the IC. Error bars represent standard error.

The plastic changes we observed in the predictable-W group replicated and extended those we reported in Cruces-Solís et al. (Cruces-Solís et al., 2018). We again found an increase in response gain and a shift in frequency tuning in this group compared to the control group. The increase in sound-evoked responses was evident when comparing the tuning curves of predictable and control animals, shown in Fig 1C across example depths of 350, 550, and 700 μm (best frequency of 16 kHz for control animals) (Fig 1C; blue *vs*. red). The increase in gain was observed across all frequency regions (Supplementary Fig 1G; blue *vs*. red). The shift in best frequency (BF) is also evident in the tuning curves and quantified in Fig 1D. In addition, we could ascertain that the upward shift, while visible across many depths beyond the exposed frequency band, did not extend throughout the inferior colliculus as we had previously assumed (Cruces-Solís et al., 2018). In fact, below 950 μm, the shift was no longer significant (two-way ANOVA, group *F*(2, 217) = 1.14, *p* = 0.3, depth *F*(6, 217) = 1.99, *p* = 0.07, interaction *F*(12, 217) = 0.51, *p* = 0.9).

The key result, however, is that the patterns were similar for animals that were predictably exposed regardless of reinforcement. Indeed, the increase in response gain and shift in BF were also present in the predictable-noW group (Fig 1C and D, Supplementary Fig 1G). While the presence of water in the corner did not imply a tight temporal association between the sound and the water, for mice only drank sporadically during a given visit and only in about two thirds of the visits (de Hoz and Nelken, 2014), the results confirm that the change in IC responses was not caused by an association between the 16 kHz contextual sound and the presence of water in the corner. Quantification of average evoked response across tone frequencies confirmed that the gain increase in both predictable-W and predictable-noW animals relative to control appeared along the IC tonotopic axis (Supplementary Fig 1G; linear mixed model [two factors: group × depth]: ctrl *vs*. noW *p* = 0.016, depth *p* < 0.0001, ctrl/pred x depth *p* = 0.02). The increase in evoked responses at greater depths is likely related to the relative shank width of the probe (A1×16-3mm-50-177) at different depths (Claverol-Tinture and Nadasdy, 2004). Nevertheless, probe positioning was consistent across different animals and cannot explain the differences between the groups. Similarly, quantification of changes in BF relative to the control across recording depths between 100 and 750 μm (the first recording, to be comparable with (Cruces-Solís et al., 2018)) also confirmed that the shift in BF was not dependent on the presence of water (Fig 1D; linear mixed effect model [two factors: group × depth]: overall group *p* = 0.03, ctrl *vs*. pred. *p* = 0, pred. *vs*. noW *p* = 0.0002, ctrl *vs*. noW *p* = 0.0005). The BF shift in the predictable-noW group, while significantly different from the control, was not as strong as in the predictable-W group, indicating that while plasticity is not dependent on the water association, its presence might strengthen it.

These results replicated and substantially extended our previous finding (Cruces-Solís et al., 2018). We confirmed that exposure to predictable sound-context associations led to an enhancement in overall response gain and a global shift in BF. Importantly, this plasticity was unsupervised since it did not depend on reinforcement.

### 2. Predictability in the contextual sounds triggers plasticity

We next tested the effect of predictability by contrasting our results with the effect of unpredictable exposure. We focused on the predictability of the sound onset, which had previously been controlled by the mouse’s entry into the corner. We trained two new groups of mice separately in the Audiobox. One group was exposed again to pure tones of 16 kHz, but this time in an unpredictable way (unpredictable-W). The other group was a control group without sound exposure as before. To make the onset of the sound unpredictable, the tone pips were activated with a variable latency between 5 to 25 seconds from visit onset and only in 80% of the visits (Fig 1E, Supplementary Fig 1A). This means that sound was not controlled by the mouse movement and was also not present for the visit’s duration. The response magnitude and BF across depths in the unpredictable-W group did not differ from those in the control group that was trained in parallel (Supplementary Fig 1B, C, H). Quantification of the average difference in BF with respect to the control group across the depths between 100 and 750 mm confirmed that there was no shift in BFs in the unpredictable-W group (Fig 1G-F; Supplementary Fig 1C inserted; linear mixed model, *p* = 0.69). Therefore, the unpredictable onset of sound exposure, coupled with a weaker sound-context association, did not elicit plasticity in the IC.

We next examined whether the absence of an effect was caused by the shorter sound-context association that resulted from a delayed sound onset, or by its unpredictability. We trained two additional groups of mice separately in the Audiobox. In both groups the sound was initiated by the mouse entering the corner but, while the predictable-W group was exposed to the predictable sound, as before, for the duration of each visit, the predictable-entry group was exposed only for the first 3 seconds (Fig 1E; Supplementary Fig 1D). The predictable-entry group showed sound-evoked responses in the IC that were comparable to those of the predictable-W group trained in parallel (Supplementary Fig 1E, F, I). The average BFs across depths were also comparable (Fig 1G-F; Supplementary Fig 1F; linear mixed model, *p* = 0.08).

When combined, the comparison across groups trained with different sound-context association patterns (Fig 1E, 2A) revealed that predictability (sound triggered by visit onset), but neither reinforcement (water in the corner) nor the exposure time length, determined the broad shift in BF in the IC (Fig 1F; the linear mixed model on 3 factors [water × onset × context], a significant effect of onset *p* < 0.0001). This shift was also visible in the tuning curves (Fig 1G).

**Fig. 2:**
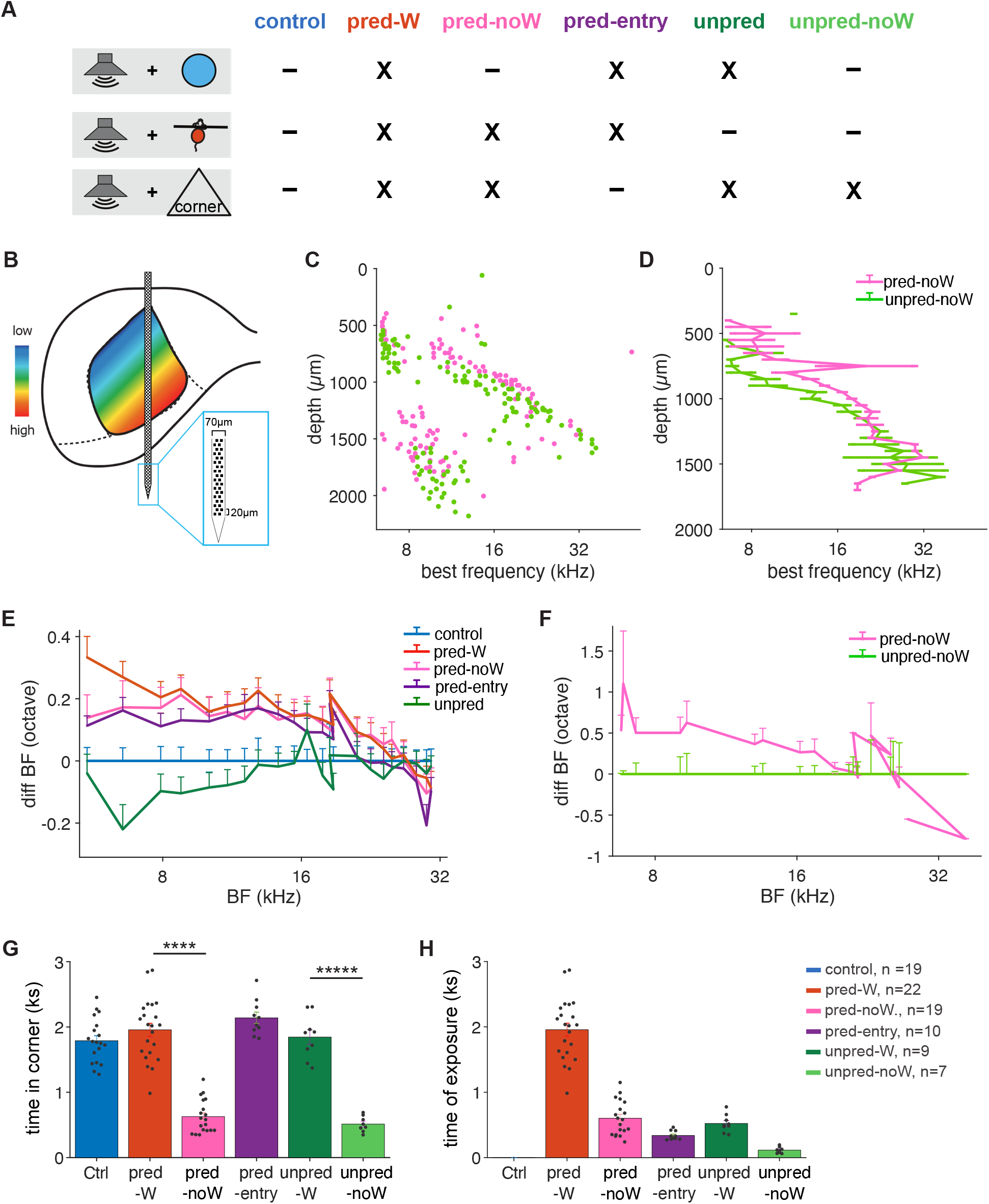
Predictability of the contextual sound triggers plasticity. **A,** Table illustrating the sound-context association pattern in different groups depending on whether there was water in the corner (top), sound triggered by visit onset (middle), sound for most of the visit (bottom). **B,** Schematic representation of recording using Neuropixels probes. Insert: Illustration of high-density Neuropixels probes. **C,** BF of individual units recorded from different depths of the IC. **D,** Average BF as a function of depths for units along the tonotopic axis. Linear mixed effect model with two factors [group × depth]: *p*_group_ = 0.016, *p*_depth_ = 0.002, *p*_group_ x depth = 0.013. **E,** Average change in BF relative to the average BF of the control group was frequency specific. **F,** Similar to **E.** for recording using Neuropixels probes. **G,** Average daily time spent in the corner of individual mice. Groups without water in the corner spent less time in the corner compared to groups with water in the corner (two-sample t-test, \*\*\*\**p*< 0.0001). **H,** Average daily length of sound exposure of individual mice.

### 3. The BF shift is broad but not ubiquitous

To better understand the frequency-specificity of the BF shift, we directly compared the effect of predictable and unpredictable sound-context associations, in the absence of water, using Neuropixels probes (Imec, Belgium) that sampled the IC in depth (Fig 2B). We trained two new groups of mice both without water in the corner, a predictable-noW and an unpredictable-noW exposure group. As expected, mice in the predictable-noW group showed an upwards shift in BF across a wide range of depths compared to the unpredictable-noW group (Fig 2C-D; linear mixed model [two factors: group × depth]: group *p* = 0.016, depth *p* = 0.002, interaction *p* = 0.013), confirming again that predictability of the sound-context association is the key variable determining the presence of plasticity. As is evident in the plots (Fig 2C-D), the shift is constrained in depth, and therefore in best frequency range.

We investigated how BF shifted along the tonotopic axis by plotting the change in BF as a function of the BF of the control group (Fig 2E). The BF shift in the predictable groups (predictable-W, predictable-noW, predictable-entry) is similar in magnitude for frequencies below 16 kHz (> 1 octave) and extends for frequencies above 16 kHz until 25 kHz (> 1/2 octave). Using a linear mixed effect model to predict the change in BF with two factors [(water + onset + context) × BF], where ‘onset’ means the sound was present upon entry, and ‘context’ means the sound could be present till the end of the visit, reveals a significant effect of onset (*p* < 0.0001) and an interaction between onset and BF (*p* < 0.0001). A similar pattern of change was observed when comparing the predictable-noW with the unpredictable-noW group (Fig 2F; linear mixed model [two factors: group × BF]: group *p* < 0.0001, BF *p* = 1, interaction *p* < 0.0001).

To sum up, we exposed six groups of mice in the Audiobox with different sound exposure paradigms and examined the effect of three possible factors: reinforcement (water in the corner), mouse entry (sound-visit onset association), and contextual value (sound present for the duration of the visit), on IC plasticity induced by sound exposure in the Audiobox (Fig 2A). Again, it is important to note that the groups were neither positively nor negatively conditioned, only exposed. Behaviorally, consistent with previous data (Cruces-Solís et al., 2018), sound exposure, independent of its duration and predictability, did not change the animals’ behavior when compared to the control group (Fig 2G-H; predictable-W, predictable-entry, unpredictable-W). These groups showed comparable daily time spent in the corner (Fig 2G, blue *vs*. red, purple and dark green; one-way ANOVA reveals no significant effect of the group, F(3,56)=2, *p* = 0.12). Interestingly, the no-water groups still visited the corner, despite the absence of water, albeit for a relatively shorter time compared to the other exposure groups (Fig 2D, red *vs*. pink, and dark green vs light green; two-sample t-test, *p* < 0.0001 for both comparisons). This resulted in a relatively short time of sound exposure (Fig 2H). What this quantification revealed is that plasticity, measured here through the magnitude of the BF shift (Fig 2E-F) could not be explained as a result of differences in the length of sound exposure, since groups with different exposure times (the predictable-entry group and the predictable-W group, for example) showed similar shift magnitudes. To conclude, sound exposure elicited a broad frequency shift in BFs along the IC tonotopic axis only when the sound onset was predicted by visit onset, and this shift was independent of the presence of a reinforcer in the corner or the length of the sound exposure (Fig 2F).

### 4. Increased frequency discriminability coupled to BF shifts

We next explored how the change in coding that was so dramatically reflected in the BF shift, affected frequency representation. First, we looked at structural population vectors: the response to a given frequency across the IC (Cruces-Solís et al., 2018), which reflect the ongoing responses to sounds in the IC. For example, when the 16 kHz sound is presented, cells at several depths in the IC will respond simultaneously with different strengths (Fig 3A, black). A different pattern will appear when a frequency farther away from 16 kHz is presented. Population vectors obtained with NeuroNexus probes, plotted side by side for each group (Fig 3B, Supplementary Fig 2B), illustrate well the increasing BF with depth characteristic of the tonotopic map of IC, but they also reveal other interesting features. For example, at greater depths, neurons often responded to low frequencies in addition to the depth-appropriate frequencies, an effect that can also be observed in the low-frequency cluster at greater depths in the Neuropixels recordings (Fig 2B), similar to the long tail of the auditory nerve (Kiang and Moxon, 1974). To quantify the dissimilarity of activity patterns between responses to different frequencies, we measured in each mouse the Euclidean distance (Edelman, 1998) between frequency-specific population vectors at different ΔF (frequency difference in octave; Fig 3C, Supplementary Fig 2C) and found increased separation in the predictable groups compared to the unpredictable group (Fig 3C and D). This effect was frequency- and ΔF-dependent (Supplementary Fig 2E-F). Modeling a linear fixed effect model to predict the boundaries where Euclidean distance is above 2 (about 80% of the max value; Fig 3D) with two factors [(water + onset + context) × frequency], reveals significant effects of frequency (*p* < 0.0001), onset (*p* = 0.04), and interactions between onset and frequency (*p* = 0.009). The dissimilarity of activity patterns not surprisingly increased with ΔF, but it did so more in the predictable groups. This increase was not limited to comparisons with the exposed frequency and, like the BF shift, extended above and below. Independently of ΔF, the discriminability in all the groups, including the control, was larger for frequencies near and above 16 kHz but weaker for frequencies below. This pattern was more patent in the predictable groups (Supplementary Fig 2F). We conclude that predictable exposure increases frequency discriminability.

**Fig. 3:**
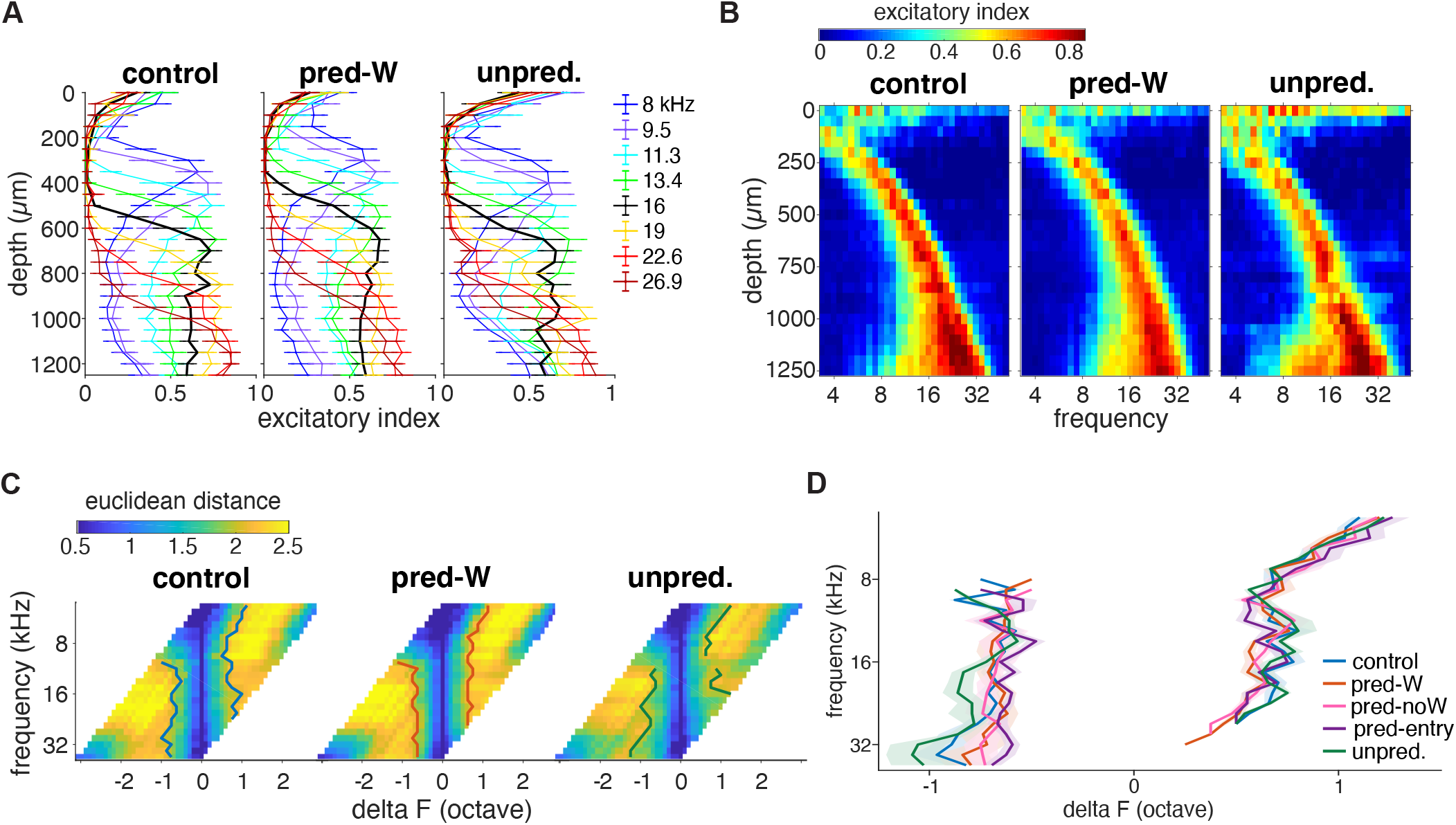
BF Shifts were accompanied by an increase in frequency discrimination ability. **A,** Population vectors, as relative response magnitude to a given frequency across the IC. Sound-evoked responses were rescaled with a 0-max normalization based on the tuning curve of each depth. **B,** Excitatory profile calculated as the relative response magnitude to a given frequency across the IC. **C,** The dissimilarity of activity patterns measured by Euclidean distance between the excitatory profile of frequency pairs. The boundaries where the mean Euclidean distance was larger than 2 were marked for each group respectively. **D**, The average boundaries across individual mice for different groups. Linear fixed effect model with two factors [(water + onset + context) × frequency]: *p*_frequency_ < 0.0001, *p*_onset_ = 0.04, *p*_onset × frequency_ = 0.009.

### 5. The BFs remained unchanged during the initial days of sound exposure

All the data shown so far were obtained from mice that had experienced context-sound associations for 6-12 days in the Audiobox. We were interested in determining whether plasticity, in the form of response gain changes and shifts in frequency representation, was observable already in the early days of exposure. For that purpose, we recorded from mice that had been in the Audiobox for only 1 to 5 days. We compared control and predictable-W groups and found that both the differences in gain and BF described earlier showed a different pattern in the early days. During the initial exposure, the control group showed generally stronger evoked activity than the predictable-W group across a wide range of depths (Fig 4A and Supplementary 3D; linear mixed model for the ventral area [two factors: group × depth]: group *p* = 0.005, depth *p* = 0.007, interaction *p* = 0.036). The BF, on the other hand, was not significantly different across depths between groups (Fig 4B). Importantly, this lack of BF shift in the early days was also observed in mice in a predictable-noW group (Supplementary 3B). Thus, the time course of plasticity was similar independently of reward. A linear mixed model [two factors: group × depth] reveals no significant effect of group (*p* < 0.05). This demonstrates that changes in response gain are independent of changes in BF shifts. For comparison, we include the late exposure mice published in Cruces-Solis et al. (2018), which were run in a parallel experiment (Fig 4C), as well as the late exposure mice from the predictable-noW and associated control groups from this study (Supplementary Figure 3C). Combining these early (1-5 days) and late (6-12 days) exposure, we confirmed that, indeed, the BF shift in the predictable-W exposed group developed over days and had different dynamics from that of the control group (Fig 4C; linear mixed model with three factors [group × training days × depth]: group *p* = 0.03, depth *p* < 0.0001, training day *p* = 0.31, group×days *p* = 0.13). Thus, the plastic changes observed in the late days (6-12) of exposure developed slowly and showed a different pattern in the early days.

**Fig. 4:**
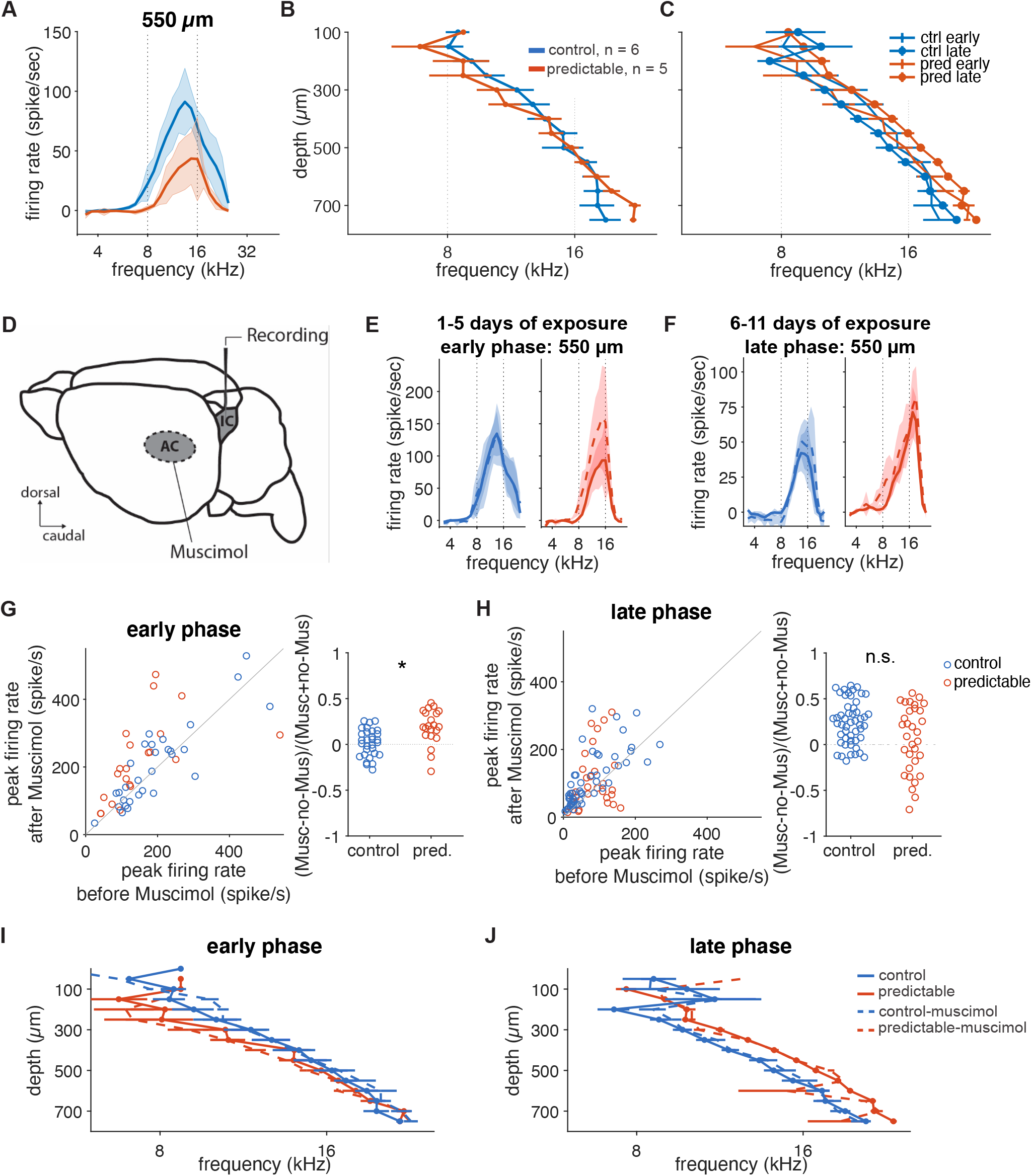
Corticofugal input suppressed collicular activity during the early days of predictable sound exposure. **A,** Average tuning curves, before (solid lines) and after cortical inactivation (dashed lines), at 550 μm for control (blue) and predictable (right) groups during the early days of sound exposure. **B**, Average best frequency (BF) as a function of recording depths for mice during the early days of sound exposure. **C**, Average best frequency (BF) as a function of recording depths for mice during the early and late days of sound exposure. **D**, Schematic representation of simultaneous inactivation of the AC and recording in the IC. **E**, Average tuning curves of recorded sound-evoked activity, before (solid lines) and after (dashed lines) muscimol application in the AC, at 70 dB for 550 μm in the IC of the control (blue) and predictable (red) group. The recording was performed from mice with no more than 5 days of sound exposure. **F**, Similar to **E**, for mice with 6 to 11 days of sound exposure. **G, Left**, pairwise comparison of peak activity before and after cortical inactivation, for mice in the early phase of sound exposure. Wilcoxon signed rank test comparing before and after cortical inactivation: control *p* = 0.30, predictable *p* = 0.003. **Right**, cortical inactivation led to stronger disinhibition in the predictable group (linear mixed effect model with the group factor: *p*group = 0.016). Animals and recording sites: control *n* = 2 and 21, predictable *n=* 3 and 30. **H**, Similar to **G**, for the late phase. Wilcoxon signed rank test comparing before and after cortical inactivation: control *p* < 0.0001, predictable *p* = 0.19. Cortical inactivation led to a comparable increase in response gain (linear mixed effect model with the group factor: *p*group = 0.24). Animals and recording sites: control *n* = 3 and 32, predictable *n=* 5 and 51. **I, J**, Average best frequency (BF) as a function of recording depths for mice during the early and late phase of sound exposure respectively.

### 6. Corticofugal input differentially suppressed collicular activity during predictable sound exposure but only during the early days

Direct cortical feedback through descending projections from deep layers of the auditory cortex (Nakamoto et al., 2013; Yudintsev et al., 2021) has been shown to induce collicular plasticity (Gao and Suga, 2000; Suga et al., 2002; Yan and Ehret, 2001), modulate collicular sensory filters (Syka and Popelár, 1984), and play a role in learning (Bajo et al., 2010; Bajo and King, 2012). We previously found that while auditory cortical inactivation (Fig 4D) led to disinhibition in the inferior colliculus, it had no differential effect on predictable and control groups exposed for 6-12 days (Cruces-Solís et al., 2018). In both, it led to a comparable increase in response gain (current Fig 4F and H) in the absence of an effect on the upward BF shift (Fig 4J). Thus, we concluded that corticofugal input modulated response gain without affecting BF shifts, and that the maintenance of the plastic changes triggered by predictable sound exposure was not dependent on cortical feedback. To test whether corticofugal inputs affect predictable learning during the initial sound exposure, we performed simultaneous inactivation of the auditory cortex with muscimol and recordings in the inferior colliculus as before, on a subset of control and predictable animals that had been exposed for only 1-5 days (see Methods, Fig 4D). Unlike in the late exposure days, the effect of cortical inactivation on IC activity was different in control and predictable groups during the early days of exposure (Fig 4E and G). Response gain was differently affected by cortical inactivation (Fig 4E and G-left; Wilcoxon signed rank test comparing before and after cortical inactivation: control *p* = 0.30, predictable *p* = 0.003.), leading to stronger disinhibition in the predictable group (Fig 4G right; linear mixed effect model with the group factor: *p* = 0.02). Regarding the BF shift, similar to during the late phase of exposure (Fig 4J), cortical inactivation had no effect on BF during the early phase of exposure (Fig 4I), supporting again the idea that BF shifts are independent of response gain changes as well as independent from cortical feedback. Thus, the auditory cortex had a stronger effect on IC responses of mice exposed to predictable context-sound associations than in control mice, but only during the early days of exposure. This suggests that the weaker responses, relative to control, that we observed in these early phases (Fig 4A and Supplementary 3D) might be due in part to cortical inhibition that is stronger in the predictable group.

## Discussion

What we hear is often predicted by where we are. Sound-context associations are informative and predictive of normality. They are learned implicitly, in an unsupervised manner. Aiming to understand better the coding of unsupervised learning of sound-context associations, we measured changes in sound evoked responses in the inferior colliculus, a hub of auditory input as well as multimodal feedback, of mice that had lived in an enriched environment, the Audiobox, and had been exposed to sound-context associations with group-specific degree of predictability over days. We found that living in an enriched environment with predictable sound-context associations led to an upward and broad frequency shift in best frequency and an increase in response gain in the inferior colliculus. These plastic changes were paralleled by increased neuronal frequency discrimination and were independent of reinforcement. Plasticity was observed in all exposure conditions in which the sound was activated as the mouse entered the corner, independently of sound exposure and independently of the presence of reinforcement. Plasticity was not observed when the sound was activated at random times after visit onset. We conclude that the auditory midbrain is sensitive to the predictability of contextual sounds and discuss the consequences this might have for the processing of background and foreground acoustic information.

The results expand previous findings from the laboratory where we found an increase in gain and, what we interpreted as a frequency-unspecific BF shift, in animals exposed to predictable sounds in the presence of water (Cruces-Solís et al., 2018). The sampling of a wider depth range with both Neuronexus and Neuropixels probes in the current study revealed that the BF shift is broad but constrained in depth. Similarly, the comparison between the effects of different forms of sound-context association in this series of experiments, in the presence and absence of reinforcement, allows us to conclude that plasticity is associated specifically with the unsupervised learning of predictable sounds. While the presence of narrow tubes in the Audiobox contributed to the natural exploration and behaviour of the mice, it made it impossible for us to perform the recordings in awake mice. Recordings were performed in anesthetized animals removed from the enriched environment at different time points from the beginning of sound exposure (early groups: 1-5 days; late groups:6 and 12 days for Neuronexus recordings and 19 to 21 days for Neuropixels). Anesthesia can affect neuronal responses (Jing et al., 2021), but it is unlikely to have different effects in different exposure groups. Our conclusions thus are drawn from the comparison across groups of mice that experienced different degrees of sound-context predictability and reflect comparative changes in sensory processing.

We have shown before that when mice are exposed to non-reinforced sound-context associations, as in the current study, they do not modify their behaviour in a sound-dependent manner (Cruces-Solís et al., 2018; de Hoz and Nelken, 2014). Also in this study, animals exposed to a predictable or an unpredictable sound, spent comparable daily time lengths in the corner as mice exposed to no sound at all. The presence or absence of water in the corner did influence the motivation to go to the corner, leading to fewer visits and shorter exposure time, but not the qualitative effects on plasticity. The lack of noticeable change in behavior after unsupervised predictable sound exposure should not be confused with a lack of learning. We have observed across many studies that non-reinforced exposure in the Audiobox leads to latent learning that is only expressed during subsequent reinforced learning (Cruces-Solís et al., 2018; de Hoz and Nelken, 2014), with different time courses than reinforced learning in the same apparatus (Chen et al., 2021, 2019; de Hoz and Nelken, 2014), and effects on reinforced associative plasticity in the auditory cortex (de Hoz et al., 2022). Unsupervised exposure does lead to unsupervised learning and we believe that the changes in midbrain auditory responses are a correlate of this learning. Indeed, we found that neuronal generalization during unsupervised exposure paralleled behavioral generalization measured through pre-pulse inhibition of the startle reflex (Cruces-Solís et al., 2018).

In the experiments described here, the sound is controlled by the mouse as it enters the corner. In the predictable groups, this entry activates the sound, whether for the length of the visit or only for 3 seconds. In the unpredictable groups, this entry is not coupled with sound onset but activates a clock that will determine the random interval after which the sound will begin. A BF shift was only observed in animals in which the sound was activated upon entry, even when only for the first 3 seconds from visit onset. While we did not test a group that activated the sound at a fixed delay upon entry, overall, the data suggest that predictability could be determined by sound changes triggered through entry into a defined environment, since this is how sound-context associations are typically defined in natural situations. Time-averaged statistics of contextual sounds, such as textures, converge within 2.5-3 seconds (McDermott et al., 2013), possibly making this the minimum length of time necessary for a contextual sound to be identified as such. We did not test shorter sounds or longer sounds with slowly converging statistics.

Frequency-specific shifts in tuning are a hallmark of auditory plasticity (Bakin and Weinberger, 1990; Blake et al., 2002; Yan et al., 2005; Zhang and Suga, 2005). Typically, when an animal is trained in a frequency conditioning paradigm, a shift in BF is observed that is localized to the population of neurons that respond to the conditioned frequency. This shift enables and parallels an increase in the representation of the relevant frequency (Blake et al., 2002). What is surprising in our results is that the shift occurs also in populations of neurons that do not respond to the exposed frequency (16 kHz) and is always an upward shift, independently of whether the BF is below or above 16 kHz. In our initial study, the shift was present throughout the wide but limited range of depths measured, making us believe that the shift might be frequency-unspecific (Cruces-Solís et al., 2018). In the current work, by sampling across a wider range of depths, we could conclude that the shift is frequency-specific but broad, extending to about ⅔ of an octave above and more than 1 octave below the exposed frequency (Fig 2D-E). Unlike inferior colliculus BF shifts triggered by reinforced learning (Zhang and Suga, 2005), the shift we observed was independent of cortical feedback at all time points. Overall, the data suggest that the mechanisms underlying this positive and broad shift are different from those observed under reinforced learning. The shift is likely tightly related to the observed changes in frequency representation in the form of increased frequency discriminability for the predictable groups. Thus, while reinforced learning leads to a local shift towards the relevant frequency, with a concomitant decrease in discriminability for nearby frequencies (Bao et al., 2013; de Villers-Sidani et al., 2007), unsupervised learning resulted in a broad upward shift that leads to overall increased frequency discriminability (this study) but wider frequency response overlap (Cruces-Solis et al., 2018).

Broad frequency shifts and increase in gain developed slowly, showing more variability during the initial days of exposure, when the control group displayed a tendency for larger responses than the predictable group. Exposure was sparse, in the sense that unlike in manually trained animals, exposure events in the form of visits to the corner could be separated by many minutes (Cruces-Solís et al., 2018). But this exposure was embedded in a rich continuous environment in which many other stimuli were present, meaning that the mice living in the Audiobox were learning more patterns than just those specifically associated with the sound-context predictability. This initial disentangling of the different patterns present in the environment is likely to engage either different circuits or the same circuits differently. For example, cortico-collicular feedback projections are known to play an important role in the development of inferior colliculus plasticity in reinforced learning (Yan and Suga, 1998) and to facilitate learning when input-response mappings change (Bajo et al., 2010). Here, while we confirm the influence of cortico-collicular projections on inferior colliculus levels of activity, what was interesting is that this cortical feedback had a differential effect on the inferior colliculus of mice exposed to predictable context-sound association compared to control mice during the initial days of exposure. This is in contrast with what happens in animals that lived in the Audiobox for at last 6 days (Cruces-Solis et al., 2018 and this study), when cortical inactivation no longer has a differential effect on the two groups. Inactivating cortex did not have an effect on the BF at any time point in neither the predictable nor the control group, demonstrating that changes in gain are independent of shifts in BF. Thus, it seems that cortico-collicular feedback is more sensitive to the specifics of the environment during the initial days of exposure and might play a more relevant role in shaping inferior colliculus activity then.

Finally, we observe these changes in the inferior colliculus, an auditory midbrain hub previously associated with short lasting (up to 60 minutes) plasticity that is dependent on cortical feedback (Gao and Suga, 2000; Yan et al., 2005). The plasticity we observe here, which developed slowly over the first 6 days and lasted up to 21 days of predictable contextual sound exposure, might have different roles. Evidence for early detection of context can be deduced from noise-invariance in cortical structures. Indeed, using naturalistic background sounds, several labs have found evidence for noise-invariant responses in cortical structures (Khalighinejad et al., 2019;

Mesgarani et al., 2014; Norman-Haignere and McDermott, 2018; Rabinowitz et al., 2012), supporting the idea that context is coded separately. One possibility is that predictable acoustic context is learned through cortico-subcortical loops and encoded in subcortical structures for the rapid processing that would facilitate noise-invariant responses higher up.

## Methods

### Animals

Female C57BL/6JOlaHsd (Janvier, France) mice were used for experiments. All mice were 5-6 weeks old (NeuroNexus experiments) or 8 weeks old (Neuropixels experiments) at the beginning of the experiment. Animals were housed in groups and in a temperature-controlled environment (21 ± 1°C) on a 12 h light/dark schedule (7am/7pm) with access to food and water *ad libitum*. All animal experiments were approved and performed in accordance with the Niedersächsisches Landesamt für Verbraucherschutz und Lebensmittelsicherheit, project license number 33.14-42502-04-10/0288 and 33.19-42502-04-11/0658; and in accordance with the Landesamt für Gesundheit und Soziales (LaGeSo), project license number G0140/19.

### Apparatus: The Audiobox

The Audiobox is a device developed for auditory research and based on the Intellicage (NewBehavior, Switzerland). Mice were kept in groups of 6 to 10 animals. At least one day before experimentation, each mouse was lightly anesthetized with Avertin i.p. (0.1ml/10g), Ketamin/Xylazin (13 mg/mL / 1 mg/mL; 0.01 mL/g), or isoflurane and a sterile transponder (PeddyMark, 12 mm × 2 mm or 8 mm × 1.4 mm ISO microchips, 0.1 gr in weight, UK) was implanted subcutaneously in the upper back. Histoacryl (B. Braun, Germany) was used to close the small hole left on the skin by the transponder injection. Thus, each animal was individually identifiable through the use of the implanted transponder. The Audiobox served both as living quarters for the mice and as their testing arena.

The Audiobox was placed in a soundproof room which was temperature regulated and kept in a 12 h dark/light schedule. The apparatus consists of three parts, a food area, a ‘corner’, and a long corridor connecting the other two parts (Fig. 1A). The food area serves as the living quarter, where the mice have access to food *ad libitum*. Water is available either in two bottles situated in the corner or in the food area. The presence of the mouse in the ‘corner’, a ‘visit’, is detected by an antenna located at the entrance of the corner. The antenna reads the unique transponder carried by each mouse as it enters the corner. A heat sensor within the corner senses the continued presence of the mouse. An antenna located at the entrance of the corner detects the transponder in the back of the mouse as it enters. The mouse identification is then used to select the correct acoustic stimulus. Once in the ‘corner’, specific behaviors (nose-poking and licking) can be detected through other sensors. All behavioral data is logged for each mouse individually. A loudspeaker (22TAF/G, Seas Prestige) is positioned right above the ‘corner’, for the presentation of the sound stimuli. During experimentation, cages and apparati were cleaned once a week by the experimenter.

### Sound exposure in the Audiobox

Sounds were generated using Matlab (Mathworks) at a sampling rate of 48 kHz and written into computer files. Intensities were calibrated for frequencies between 1 and 18 kHz with a Brüel & Kjær (4939 ¼” free field) microphone.

All mice had *ab libitum* food in the home cage, as well as free access to water through nosepoking in the corner, except the ‘no-water’ groups (noW; see below) which had water in the food area instead. There was no conditioning in the Audiobox at any stage. All groups were first habituated to the Audiobox for 3 days without sound presentation. After the habituation phase, except for the control group, mice were exposed to 16 kHz pure tone pips in the corner with different exposure protocols.

The ‘predictable-W’ group had the same sound exposure paradigm as in a previous study (Cruces-Solís et al., 2018). The mice were exposed to 16 kHz tone pips for the duration of each visit, regardless of nose-poke activity and water intake.

The ‘predictable-noW’ group was treated the same as the predictable-W group, i.e., it was exposed to 16 kHz tone pips for the duration of each visit. However, there was no water in the corner, and water was accessible in the home cage.

For the ‘predictable-entry’ group, 16 kHz tone pips were triggered by the entry into the corner, but only lasted for 3 sec. For visits with a duration of less than 3 sec, the sound stopped once the mouse left the corner.

For the ‘unpredictable’ group, 16 kHz tone pips were presented in 80% of the visits with delay random between 5 to 25 seconds from the visit onset. Once the sound started, it continued until the mouse left the corner. Thus, the onset of the sound could not be predicted by the animal.

The control group lived in the Audiobox for the same amount of time as the other groups, but without sound presentation. All animals can hear environmental sounds, i.e., their own vocalization and the sound of the ventilation fan. All animals were in the exposure phase for at least 5 days. All animals were in the Audiobox in the exposure phase for at least 5 days: 6-12 days in the Neuronexus data set and 19-21 days in the Neuropixels data set.

### Electrophysiology

After at least 5 days in the exposure phase, animals were removed from the apparatus one at a time and immediately anesthetized for electrophysiology. Animals were initially anesthetized with Avertin (0.15ml/10 g; Neuronexus recordings) or Ketamine/Xylazine (13 mg/mL / 1 mg/mL; 0.01 mL/g; Neuropixels recordings). Additional smaller doses of drug were delivered as needed to maintain anesthesia during surgery. The surgical level of anesthesia was verified by the pedal-withdrawal reflex. Body temperature was maintained at 36 °C with a feedback-regulated heating pad (ATC 1000, WPI, Germany).

Anesthetized mice were fixed with blunt ear bars on a stereotaxic apparatus (Kopf, Germany). Vidisic eye gel (Bausch + Lomb GmbH, Germany) was used to protect the eyes from drying out. Lidocaine (0.1 mL) was subcutaneously injected at the scalp to additionally numb the region before incision. The scalp was removed to expose the skull. Periosteum connective tissue that adheres to the skull was removed with a scalpel. The bone surface was then disinfected and cleaned with hydrogen peroxide. Bone suture junctions Bregma and Lamda were aligned. A metal head-holder was glued to the skull 1.0 mm rostral to Lambda with methyl methacrylate resin (Unifast TRAD, GC), or a combination of Histoacryl (B. Braun) and dental cement (Lang Dental and Kulzer). A craniotomy of 0.8 mm × 1.0 mm with the center 0.85 mm to the midline and 0.75 caudal to Lambda was made to expose the left inferior colliculus. After the craniotomy, the brain was protected with Saline (B. Braun, Germany). The area which is posterior to the transverse sinus and anterior to the sigmoid sinus was identified as the inferior colliculus.

The borders of the left inferior colliculus became visible after the craniotomy. The electrode was placed in the center of the inferior colliculus after measuring its visible rostrocaudal and mediolateral borders. The aim was to target the same position of the inferior colliculus across animals. Before lowering the electrode, a ground wire was connected to the neck muscle as a reference for the recordings. Neuronexus electrodes (single shank 16-channel silicon probe; 177 μm^2^ area/site and 50 μm spacing between sites; Neuronexus, USA) were vertically lowered with a micromanipulator (Kopf, Germany) and slowly advanced (2-4 μm/sec, to minimize damage to the tissue) to a depth of 750 μm from the brain surface. The final depth was double-checked by making sure that the most dorsal channel was aligned with the dura (Fig. 1B). After one recording session, the electrode was further advanced 500 μm to a depth of 1250 μm. A final set of experiments was performed in Berlin using Neuropixels probes (Imec, Belgium, 1.0 with cap). These were inserted 3000 μm deep into the central nucleus of IC measured from the surface of the dura, at a speed of 62 μm per second using an electronic manipulator (Luigs & Neumann). A silver ground was attached to the probe and positioned at the caudal end of the skin incision.

During data acquisition with Neuronexus probes, the electrophysiological signal was amplified (HS-18-MM, Neuralynx, USA), sent to an acquisition board (Digital Lynx 4SX, Neuralynx, USA), and recorded with a Cheetah 32 Channel System (Neuralynx, USA). The voltage traces were acquired at a 32 kHz sampling rate with a wide band-pass filter (0.1±9,000 Hz). Neuropixels recordings were performed using a PXIe-6341 board (National instrument), sampled at 30 kHz, and recorded with open-source software SpikeGLX (http://billkarsh.github.io/SpikeGLX/).

### Acoustic stimulation during recording

The sound was synthesized using MATLAB, produced by a USB interphase (Octa capture, Roland, USA), amplified (Portable Ultrasonic Power Amplifier, Avisoft, Germany), and played with a free-field ultrasonic speaker (Ultrasonic Dynamic Speaker Vifa, Avisoft, Germany). The speaker was 15 cm away from the right ear. The sound intensity was calibrated with a Bruël & Kjaer microphone. For measuring the frequency response area, we used sound stimuli which consisted of 30 ms pure tone pips with 5 ms rise/fall slopes repeated at a rate of 2 Hz. Thirty-two frequencies were used (3.4 kHz to 49.4 kHz, 0.125-octave spacing) at different intensities (0 dB to 80 dB with steps of 5 or 10 dB) played in a pseudorandom order. Each frequency-intensity combination was played 5 times. For Neuropixels recordings, 51 frequencies were used (6.5 to 65 kHz), played at 70 dB SPL, each repeated 5 times in a given sweep. Two or three sweeps were played per recording.

### Analysis of electrophysiological recordings

The recorded voltage signals were high-pass filtered at 500 Hz to remove the slow local field potential signal. To improve the signal-to-noise ratio of the recording, the common average reference was calculated from all the functional channels and subtracted from each channel (Ludwig et al., 2009).

For Neuronexus data multi-unit analysis, spikes were detected as local minima below a threshold of 6 times the median absolute deviation of each channel and later summed at 1ms bins. If the calculated value was higher than −40 μV, the threshold was set to −40 μV. Multi-unit sites were included in the analysis only if they were classified as sound-driven, defined by significant excitatory evoked responses (comparison of overall firing rate in 80 ms windows before and after stimulus onset for all frequencies and intensities; paired t-test, *p* < 0.05).

To analyze the sound-driven responses, signals were aligned at 200 ms before each stimulus onset. The temporal pattern of responses was compared using peri-stimulus time histograms (PSTHs). Multiunit spikes were averaged over an 80 ms window before and after each stimulus onset to calculate the spontaneous activity and evoked firing rates respectively.

Sound-driven responses at a given intensity as a function of frequency were used to generate the iso-intensity tuning curves. The combined frequency-intensity responses were used to generate the tonal receptive field (TRF).

The best frequency (BF) was selected as the frequency which elicited the highest responses when averaging across all intensities. In the rare cases where more than one frequency elicited the highest response, BF was calculated as the mean of those frequencies.

The excitation profile was characterized by computing the normalized response to a given frequency across different depths. The normalization was done by rescaling the tuning curve of each depth with a 0-max normalization. Discriminability between frequency pairs was calculated as Euclidean distance between their population vectors.

Neural activities at the depth from 500 μm to 750 μm were measured in two recording sessions, first when the probe tip was at 750 μm depth and then when it was at 1250 μm. When analyzing responses at a given depth, recordings for the same depth were averaged.

Spike trains from the Neuropixels data set were sorted via the open-source platforms Kilosort3 and CatGT. Frequency tuning was derived using custom scripts built in MATLAB 2020a. Evoked activity was calculated as all tuned post-trigger spikes summed over an 80 ms time window. Only activity which was significantly different from an 80 ms pre-trigger window was considered evoked. A set of selection criteria were applied to these evoked units: 1) we imposed a minimum spike amplitude ≥ 18 μV, which corresponds to 3 times the baseline noise (Steinmetz et al., 2018); 2) a minimum spike count of 25, across responses to all frequencies within a frequency sweep; 3) tuning, meaning a mooth curve with no more than 4 crossings of the median spike count line.

To remove the tail units from Neuropixels datasets in order to construct the mean responses plotted in Figure 3C, a two-cluster Gaussian mixture model (GMM; *fitgmdist*, MATLAB) initialized with the k-means algorithm was applied.

### Simultaneous cortical inactivation and recordings in the inferior colliculus

After the craniotomy to expose the inferior colliculus, another 4×3 mm rectangular craniotomy medial to squamosal suture and rostral of the lambdoid suture was performed over the left hemisphere to expose the ipsilateral AC (Fig 4A). Vaseline was applied to the boundaries of the craniotomy to form a well. Initially, recording in the inferior colliculus was performed while 3-5μL phosphate-buffered saline solution (B. Braun, Germany) was applied to the AC every 15 min. Then, saline was removed from the well over the AC with a sterile sponge. 3-5 μL muscimol (1 mg/mL, dissolved in phosphate-buffered saline solution) was applied over the AC every 15 min. Previously, we found that the muscimol application inactivated the AC usually in 15-20 min (Cruces-Solís et al., 2018). Therefore, we recorded the neuronal activity of the IC again 20 min after muscimol application.

### Statistical analysis

Group comparisons were made using multiple-way ANOVAs after testing for normality distribution using the Shapiro-Wilk test. Samples that failed the normality test were compared using a Kruskal-Wallis test or Wilcoxon signed rank test for paired datasets. Multiple comparisons were adjusted by Bonferroni correction. For the analysis of data consisting of two groups, we used either paired t-tests for within-subject repeated measurements or unpaired t-tests otherwise. For data consisting of more than two groups or multiple parameters we used, repeated-measures ANOVA. All multiple comparisons used critical values from a t distribution, adjusted by Bonferroni correction with an alpha level set to 0.05.

Certain analyses were performed using linear mixed-effects models (*fitlme*, MATLAB), as these models are appropriate for data with repeated measurements from the same subject. The mouse was used as the random effect. We checked whether the residual of the linear mixed-effects model was normally distributed. A significant test was performed by comparing the full model with a similar model with only random effects using the standard likelihood ratio test.

Means are expressed ± SEM. Statistical significance was considered if *p* < 0.05.

## Acknowledgements

We are grateful to Markus Krohn, head of the technical workshop at the MPI-EM, and his team, to Harry Scherer, and to the Charite technical workshop for technical support.

CC is funded by Max Plank Institute Ph.D. scholarship; AE was funded by Einstein grant EVF-2021-618.

## Author Contributions

CC and LDH conceptualized and designed the study. CC, HC and AE were involved in data collection and processing. CC analyzed the data and prepared the figures. CC and LDH wrote the manuscript, and all coauthors revised and approved it.

## Figure legends

**Supplementary Fig. 1: Predictable sound-context association induced a homogeneous increase in response gain.**

**A,** Schematic representation of the sound exposure protocol for the control and unpredictable group. **B,** Average tuning curves of simultaneously recorded sound-evoked activity at 70 dB for the depth of 350, 550, and 700 μm in the IC of the control (blue) and unpredictable (dark green) group. **C,** Average best frequency (BF) as a function of recording depths (linear mixed effect model with the group and depth factor: *p*_group_ = 0.63, *p*_depth_ < 0.0001, *p*_group, depth_ = 0.64). Insert: Difference in BF relative to the average BF of the control group (two-sample t-test, *p* = 0.75). **D, E, F,** Similar to **A, B, C,** for the comparison between the predictable and entry-only group. There was no shift in BF (linear mixed effect model with the group and depth factor: *p*group = 0.39, *p*depth < 0.0001, *p*group, depth = 0.04) and no significant difference in BF change (two-sample t-test, *p* = 0.81). **G,** Average evoked responses to frequencies at 70 dB as a function of recording depth (linear mixed effect model with the group and depth factor: *p*_ctrl-noW_ = 0.02, *p*depth < 0.0001, *p*pred, depth ^=^ 0.02). **H,I**, Similar to **G,** show the response gain for the control vs. unpredictable (linear mixed effect model with the group and depth factor: *p*_group_ = 0.53, *p*_depth_ < 0.0001, *p*_group_, depth = 0.72) and predictable vs. entry-only (linear mixed effect model with the group and depth factor: *p*_group_ = 0.69, *p*_depth_ < 0.0001, *p*_group, depth_ = 0.006), respectively.

**Supplementary Fig. 2: BF Shifts were accompanied by changes in sound representations.**

**A,** Structural tuning curves, as relative response magnitude to a given frequency across the IC. Sound-evoked responses were rescaled with a 0-max normalization based on the tuning curve of each depth. **B,** Population vector calculated as the relative response magnitude to a given frequency across the IC. **C,** The dissimilarity of activity patterns measured by Euclidean distance between the excitatory profile of frequency pairs. The boundaries where the mean Euclidean distance was larger than 2 were marked for each group respectively. **D,** Frequency discrimination ability measured by Euclidean distance between the excitatory profile of frequency pairs. **E,** Frequency discrimination ability for a given frequency to other frequencies. **F,** frequency discrimination ability for a given ΔF as a function of sound frequency.

**Supplementary Fig. 3: The BFs remained unchanged during the initial days of sound exposure.**

**A,** Average tuning curves, before (solid lines) and after cortical inactivation (dashed lines), at 550 μm for control (blue) and noW exposure (pink) groups during the early days of sound exposure. **B**, Average best frequency (BF) as a function of recording depths for mice during the early days of sound exposure. **C**, Average best frequency (BF) as a function of recording depths for mice during the early and late days of sound exposure. **D**, Average evoked responses to frequencies at 70 dB as a function of recording depth for the predictable (red) and the control (blue) group (linear mixed effect model, for the ventral area, with the group and depth factor: *p*_group_ = 0.005, *p*_depth_ = 0.007, *p*_group, depth_ = 0.036. **E**, Similar to **D**, for the noW exposure (pink) and the control (blue) group. Modeling with a linear mixed model reveals a significant effect of the depth *p* < 0.0001, but no effect of the group *p* = 0.08. **F**, Difference in BF relative to the average BF of the control group. There is no significant difference between groups (unpaired t-test, *p* > 0.05).

