## Supplementary figures for "Subcortical coding of predictable and unsupervised sound-context associations"

### Supplementary Fig. 1

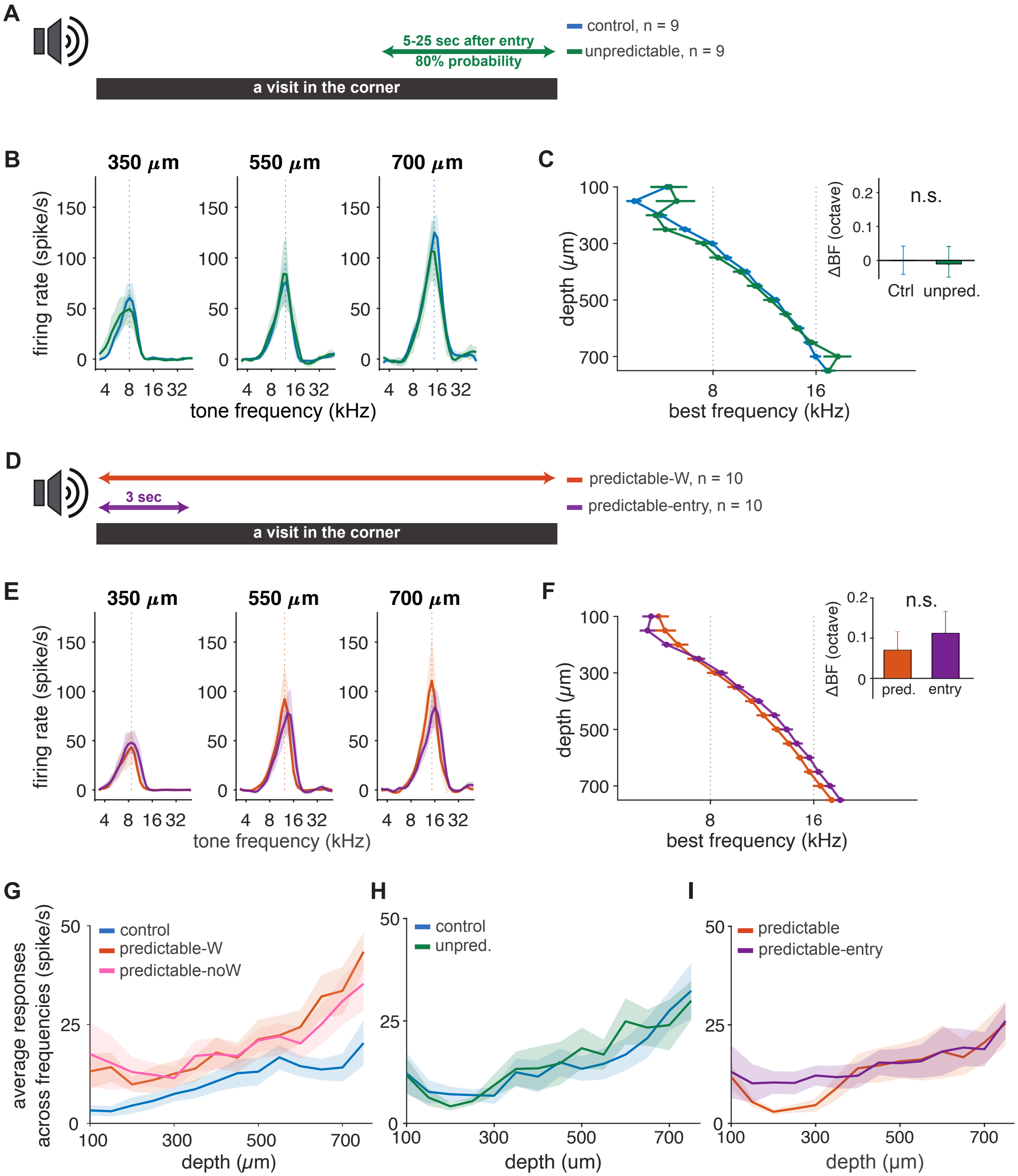

### Supplementary Fig. 2

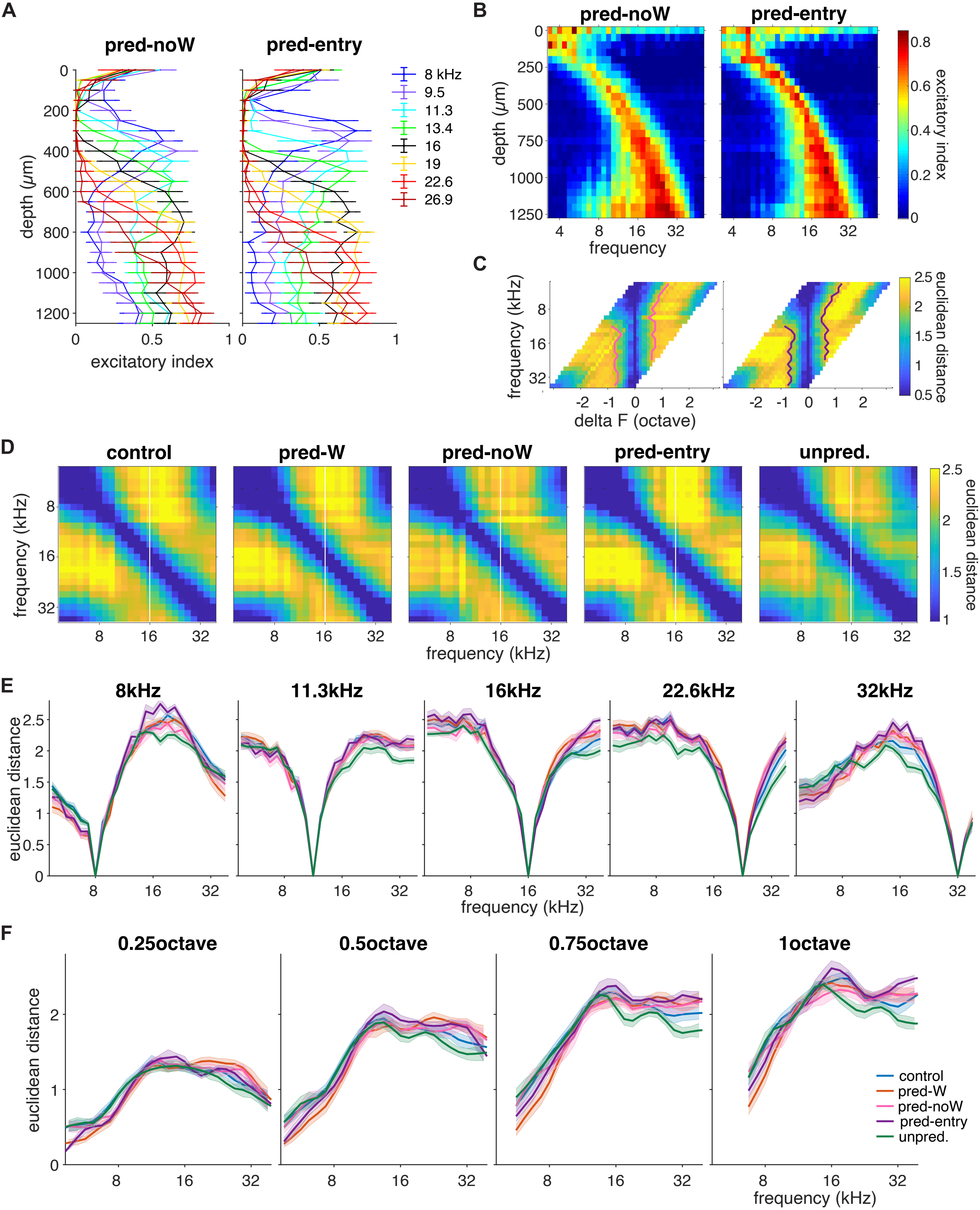

Supplementary Fig. 3

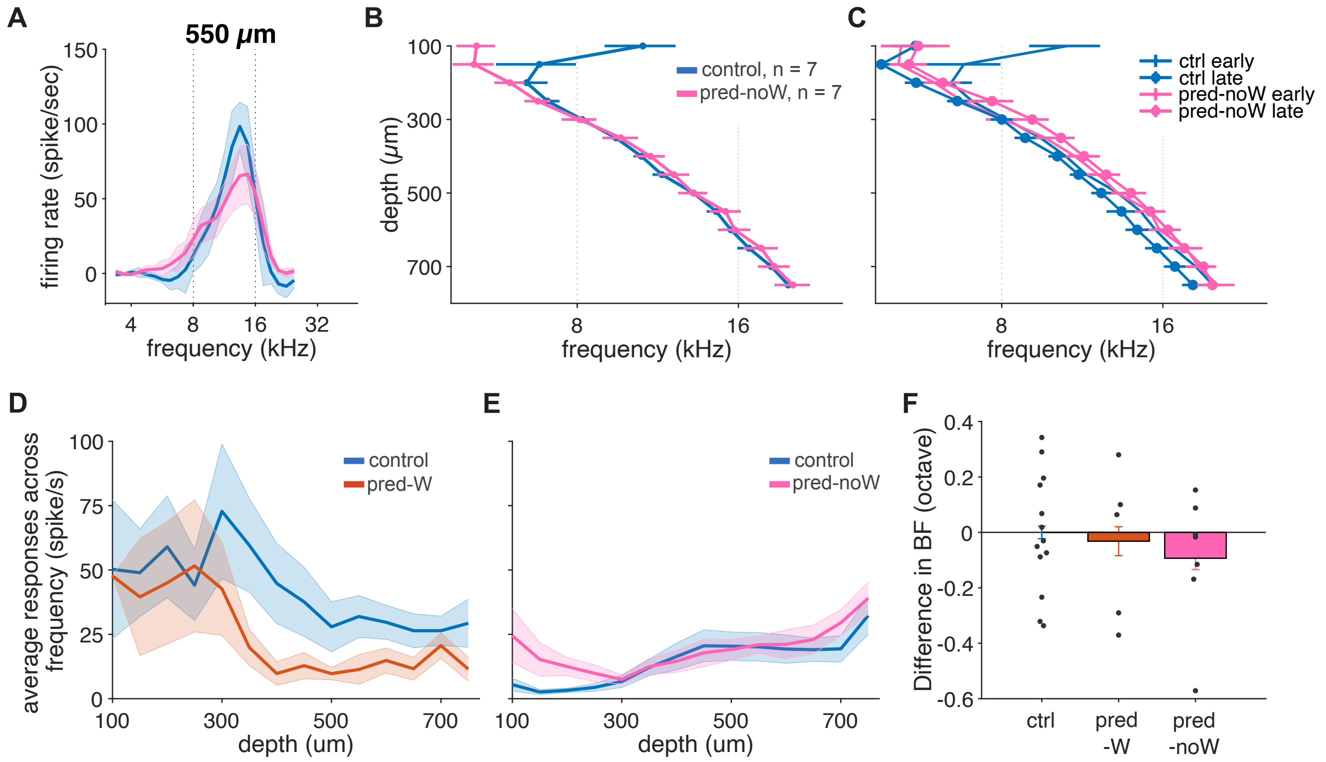

**Supplementary Fig. 1: Predictable sound-context association induced a homogeneous increase in response gain.**

**A,** Schematic representation of the sound exposure protocol for the control and unpredictable group. **B,** Average tuning curves of simultaneously recorded sound-evoked activity at 70 dB for the depth of 350, 550, and 700 μm in the IC of the control (blue) and unpredictable (dark green) group. **C,** Average best frequency (BF) as a function of recording depths (linear mixed effect model with the group and depth factor: $p_{group}$ = 0.63, $p_{depth}$ < 0.0001, $p_{group, depth}$ = 0.64). Insert: Difference in BF relative to the average BF of the control group (two-sample t-test, $p$ = 0.75). **D, E, F,** Similar to **A, B, C,** for the comparison between the predictable and entry-only group. There was no shift in BF (linear mixed effect model with the group and depth factor: $p_{group}$ = 0.39, $p_{depth}$ < 0.0001, $p_{group, depth}$ = 0.04) and no significant difference in BF change (two-sample t-test, $p$ = 0.81). **G,** Average evoked responses to frequencies at 70 dB as a function of recording depth (linear mixed effect model with the group and depth factor: $p_{ctrl\text{-}noW}$ = 0.02, $p_{depth}$ < 0.0001, $p_{pred, depth}$ = 0.02). **H,I,**Similar to **G,** show the response gain for the control vs. unpredictable (linear mixed effect model with the group and depth factor: $p_{group}$ = 0.53, $p_{depth}$ < 0.0001, $p_{group, depth}$ = 0.72) and predictable vs. entry-only (linear mixed effect model with the group and depth factor: $p_{group}$ = 0.69, $p_{depth}$ < 0.0001, $p_{group, depth}$ = 0.006), respectively.
